# Kre6-Dependent β-1,6-glucan Biosynthesis Only Occurs in the Conidium of *Aspergillus fumigatus*

**DOI:** 10.1101/2025.05.29.656687

**Authors:** Kalpana Singh, Christine Henry, Anne Beauvais, Yifan Xu, Isabelle Mouyna, Jean-Paul Latgé, Tuo Wang

## Abstract

The structural role of β-1,6-glucan has remained under-investigated in filamentous fungi compared to other fungal cell wall polymers, and previous studies have shown that the cell wall of the mycelium of *A. fumigatus* did not contain β-1,6-glucans. In contrast, the current solid-state NMR investigations showed that the conidial cell wall contained a low amount of β-1,6-glucan. ssNMR comparisons of the *A. fumigatus* and *C. albicans* β-1,6-glucans showed they are structurally similar. Deletion of the *KRE6* gene which is the only *KRE* gene in the *A. fumigatus* genome resulted in a mutant depleted of β-1,6-glucan which has a growth phenotype similar to the parental strain. Even though it is not an essential polymer in *A. fumigatus*, β-1,6-glucan play a role in cell wall organization since the *kre6*Δ mutant showed a higher sensitivity to Congo-red and Calcofluor white which are known to be general cell wall inhibitors. It is also another example of the significant structural differences seen between conidium and mycelium of filamentous fungi.

## INTRODUCTION

The cell wall of *Saccharomyces cerevisiae* and other yeasts contains two types of β-glucans. In the yeast, branched β-1,3-glucan accounts for ~30-35%, whereas β-1,6-glucan also represents 30-35% of the total yeast cell wall polysaccharides (1, 2). The *KRE* gene family (14 members) and their homologues, *SKN1* and *KNH1*, have been reported to be involved in β-1,6-glucan synthesis in yeasts (3). Among all of these genes, the ones that seem to play the major synthetic role are *KRE5* and *KRE9*, since their disruption caused a major reduction in the cell wall β-1,6- glucan content, with reductions of 100 and 80% respectively when compared to the wild type (4). Filamentous fungi, especially medically important fungi contain also β-1,6-glucan. However, the biosynthesis of β-1,6-glucan has not been precisely investigated in these fungi. In *Aspergillus fumigatus*, no β-1,6-glucan was found in the mycelium of this species (5, 6). Nevertheless, a *KRE6* ortholog was present in the *A. fumigatus* genome. The intriguing presence of a *KRE6* ortholog in the *A. fumigatus* genome in absence of β-1,6-glucan in the mycelial cell wall leads us to revisit the occurrence of β-1,6 glucans in *A. fumigatus* and characterize the function of the Kre6 protein in this fungus.

To investigate the chemical nature and molecular composition of the conidia of wild-type *A. fumigatus* and its *kre6*Δ mutant, we measured multidimensional ^13^C solid-state NMR experiments using cross-polarization (CP)-based transfer to enhance signals arising from rigid molecules. Recently, solid-state NMR analysis of the mycelial cell wall of *A. fumigatus* has suggested that α-1,3-glucans are spatially packed with chitin and β-1,3-glucan, and these carbohydrate cores re likely distributed in a soft and hydrated matrix formed by diversly linked β-glucans (6-8). Another solid-state NMR study of conidial cell walls of *A. fumigatus* has shown the reorganization of chitin, α-1,3-glucan and β-1,3-glucan during the germination process (9). In this work, we coupled NMR studies with a functional genomic approach using◻*A. fumigatus kre6*Δ mutants to identify specific structural and compositional changes associated with the absence of this gene.

## RESULTS

### β-1,6-glucans are only present in the conidia of *A. fumigatus*

To investigate the structural organization of the *A. fumigatus* cell wall, we compared the conidial and mycelial forms in the *akuB*^*ku80*^ strain. The 2D ^13^C-^13^C 53-ms CORD correlation spectra (10, 11) allowed for resonance assignment by tracing the carbon connectivity within each individual carbohydrate unit using the CORD spectra and by comparing the observed chemical shifts (**Table S1**) with previously reported values (6, 12-14). The spectra revealed a similar set of rigid polysaccharides in both morphotypes, including α-1,3-glucan, β-1,3-glucan and chitin (**Fig. 1A**). Notably, β-1,6-glucan signals were detected exclusively in the conidial cell wall, indicating a developmental regulation of this polysaccharide. These spectroscopic observations were numerically presented as the molar composition (**Table S2**), determined by averaging the peak volume of resolved signals within each monosaccharide type and the polysaccharide formed by these units. *A. fumigatus akuB*^*ku80*^ contained approximately 80 % of β-1,3-glucans, 10% of chitin, 2% of α-glucans and 8% of β-1,6-glucan.

**Fig 1.**
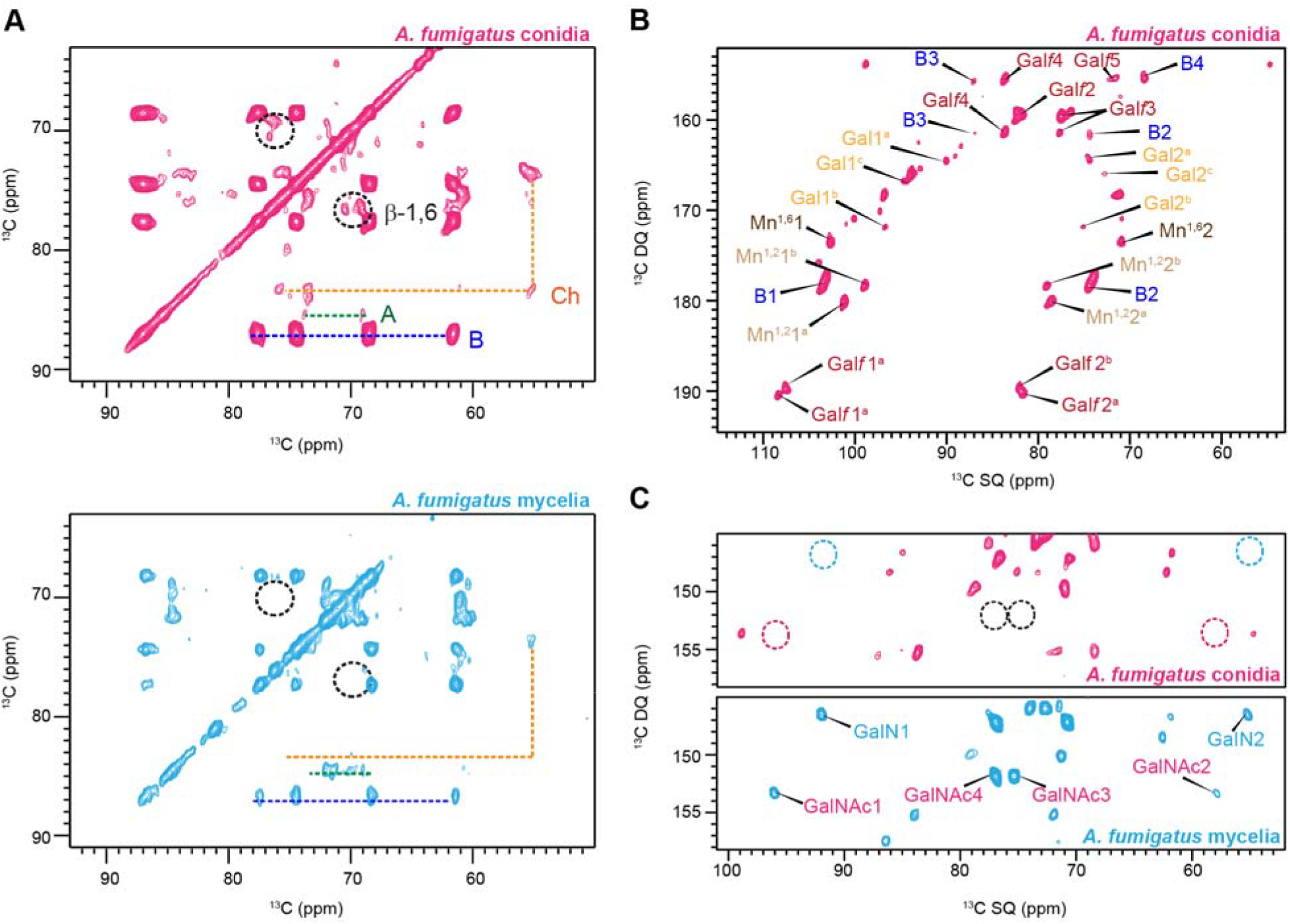
Molecular differences between *Aspergillus* conidia and mycelia. (**A**) 2D ^13^C-^13^C 53-ms CORD spectra of *A. fumigatus* conidia (magenta) and mycelia (cyan). This experiment selectively detects rigid polysaccharides. β-1,6-glucan is only present in conidia as a rigid polysaccharide. NMR abbreviations are used. Chitin: Ch; β-1,3-glucan: B; α-1,3-glucan: A. (**B**) Mobile components detected by 2D ^13^C DP refocused *J*-INADEQUATE spectra of *A. fumigatus* conidia. Mn^1,2^: α-1,2-linked Man; Mn^1,6^: α-1,6-linked Man. (**C**) Zoomed view of the specific spectral region for galactosaminogalactan, where the GalN and GalNAc units are detected in mycelia but not in conidia.

The mobile fraction was analyzed using a refocused J-INADEQUATE experiment coupled with a short recycle delay of 2 second, which select dynamic components with fast ^13^C longitudinal relaxation (**Fig. 1B**). In both conidia and mycelia, signals corresponding to α-1,2- and α-1,6- linked mannose residue (Mn^1,2^ and Mn^1,6^), which constitute the galactomannan (GM) backbone were detected (6, 15, 16). Additionally, the galactofuranose (Gal*f*) residue forming the GM side chains were observed in both forms (17, 18). Intensity analysis showed that the mobile fraction of *A. fumigatus akuB*^*ku80*^ conidia cell wall primarily consists of GM (68%) with a smaller amount of β-1,3-glucans (32%; **Table S3**). The observations of GM, along with presence of β- 1,3-glucans, support their prominent roles in the mobile domains of cell walls, which encompass the outer surface and the soft matrix. Meanwhile, it is important to note that β-1,6-glucan was absent from the mobile fraction of both mycelial and conidial cell walls.

As in the rigid region, the mobile polysaccharide fraction showed significant differences between conidia and mycelia. Galactosamine (GalN) and N-acetylgalactosamine (GalNAc), two of the three monosaccharide units (alongside Gal*p*) that compose the heteroglycan galactosaminogalactan (GAG), were absent in conidia. (**Fig. 1C**) (19). The presence of GalNAc and GalN residues in GAG was confirmed in mycelia by their characteristic single quantum (SQ) chemical shifts at 55 and 57 ppm (C2) which was further validated by correlations to C1 resonance at 92 and 96 ppm with corresponding double-quantum (DQ) shifts at 147 and 153 ppm (**Fig. 1C**). Overall, the cell walls of *A. fumigatus* mycelia and conidia exhibit distinct compositional and structural differences that reflect their specialized biological roles.

### NMR and genomic data confirm the exclusive presence of β-1,6-glucan in conidial cell wall

Recently, we analyzed the β-1,6-glucan within the mobile region of the *Candida* cell wall and successfully traced the complete NMR signals of this molecule (14). To confirm the presence of β-1,6-glucans in *A. fumigatus*, we overlaid the sheared DP-INADEQUATE spectrum of *Candida albicans*, which detects the mobile fraction where β-1,6-glucan showed strong signals, with the CORD spectrum of *A. fumigatus* conidia, which detects the rigid fraction where putative signals of β-1,6-glucan were observed (**Fig. 2**). This comparison unambiguously resolved the carbon connectivity corresponding to β-1,6-glucan in the rigid region of *A. fumigatus* conidial cell wall.

**Fig 2.**
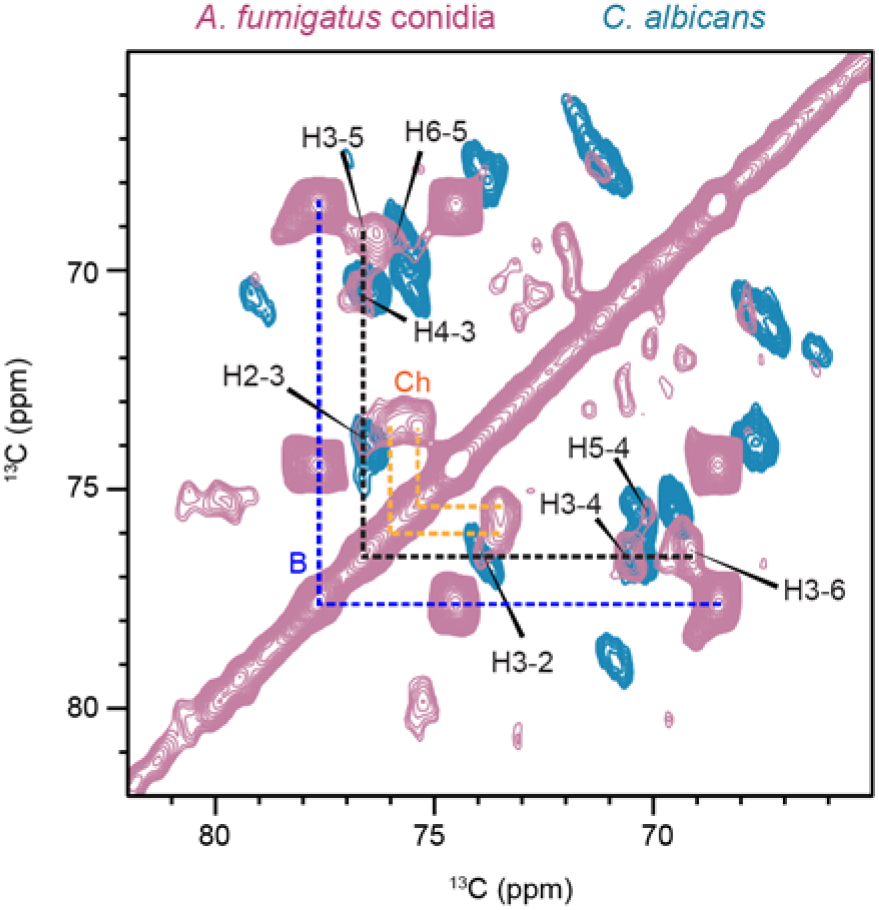
Overlay of ^13^C-^13^C CORD spectrum of *A. fumigatus akuB*^*ku80*^ (pale pink) with the sheared DP J-INADEQUATE spectrum of *C. albicans* (blue). Dashed lines highlight the major carbon connectivity for β-1,6-glucan (black), β-1,3- glucan (blue) and chitin (orange). NMR abbreviations are used. H: β-1,6-glucan; B: β- 1,3-glucan; Ch: chitin.

*KRE* genes and in particular *KRE5,6* and 9 genes are known to play an essential role in β-1,6- glucans synthesis in yeast (2). Kre5 and Kre9 proteins have no orthologs in *A. fumigatus* and a single *KRE* gene, *KRE6* has been found in the genome of *A. fumigatus*. This gene (AFUA_2G11870) showed 50% identity with *S. cerevisiae* and *C. albicans KRE6* sequences (**Fig. S1**). The sequences of *A. fumigatus* and yeast *KRE6 genes* showed comparable putative UDP glucose binding sites: R^420^VSG in *S. cerevisiae, N*^444^VTG in *C. albicans* and D^363^VSG in *A. fumigatus*. A targeted deletion of the *KRE6* gene in *A. fumigatus* was performed to elucidate its role in β-1,6-glucan biosynthesis and cell wall architecture (**Fig. S2** and **Table S4**). The parental strain which is the 1163*akuB*^KU80^ and two transformants with a deletion of the *KRE6* genes were analyzed. Both rigid fractions of the parental strain and *kre6*Δ mutants were composed of similar concentrations of β-1,3-glucans, chitin and α-glucans (**Fig. 3A, B**). β-1,6-glucan was absent in the *kre6*Δ mutant and was partially replaced by β-1,3/1,4-glucan, a well-known alternated glucan previously identified (**Fig. 3C** and **Fig. S3**). Within the mobile fraction of *aku*^*ku80*^ strain and *kre6*Δ mutants, the GM content and β-1,3-glucan levels were comparable, with values of 55% and 59% for GM content, and 45% and 41% for β-1,3-glucans, respectively (**Fig. S4-S6**). This indicates that the deletion of *KRE6* did not alter the composition of the mobile fraction.

**Fig 3.**
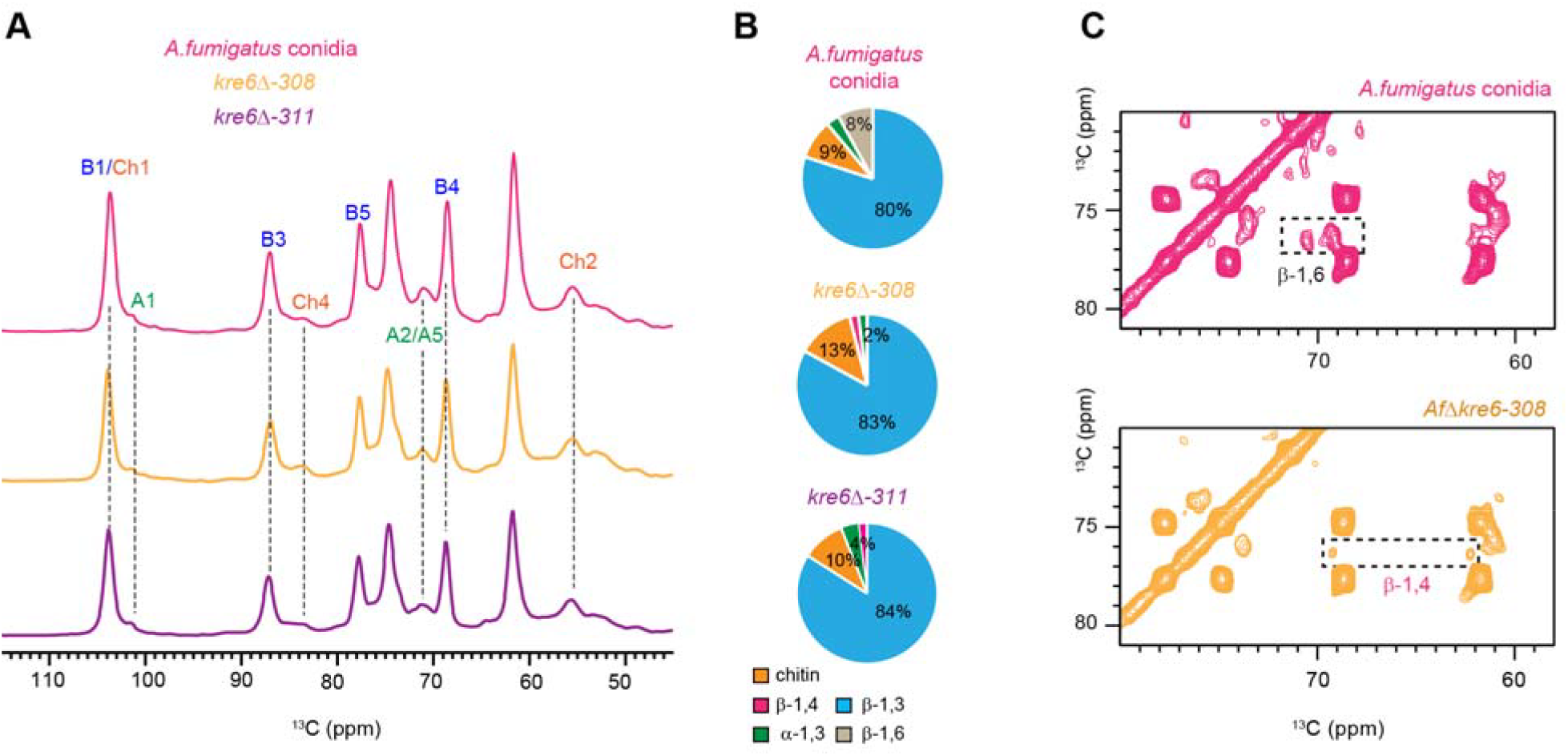
Comparison of the rigid carbohydrate components in the parental and *kre6*Δ mutants conidial cell walls. (**A**) 1D ^13^C CP spectra for selecting rigid carbohydrates showing the spectra of *A. fumigatus akuB*^*ku80*^ (magenta), *kre6*Δ mutants (*kre6*Δ-*308*, yellow; *kre6*Δ-*311*, purple). (**B**) Molar composition of the rigid polysaccharides (chitin, β-1,4-Glc units in β-1,3/1,4-glucan, β- 1,3-glucan, β-1,6-glucan, and α-1,3-glucan) in *A. fumigatus akuB*^*ku80*^ conidial cell walls and *kre*6Δ mutants determined by peak volumes of 2D ^13^C-^13^C CORD spectrum. (**C**) 2D ^13^C-^13^C CORD of *kre6*Δ mutant *Kre6*Δ-*308* (yellow) with zoomed spectra showing β-1,6-glucan in the parental strain (magenta) and the presence of β-1,4-Glc units in the β-1,3/1,4-glucan in *kre6*Δ mutant.

The phenotype of the *kre6*Δ mutant was characterized. The conidiation of the *kre6*Δ mutant was similar to the parental strain *akuB*^*ku80*^. Germination kinetics of the wild type and the mutant were similar, with 100% of the conidia germinated after 8 h incubation at 37°C on Sabouraud plates. Total growth in Sabouraud medium shaken flasks at 150 rpm 37°C, estimated as mycelia dry weight, was not affected by the gene deletion. No differential growth phenotypes were observed after incubation of the mutant at different temperatures (37°C, 45°C or 50°C) or on different media (Malt, RPMI, minimal medium, Sabouraud, supplemented or not by NaCl (1.5M) (data not shown). In our experimental conditions, which included growth in minimal medium at 37°C for 48 hours(20), resistance of *kre6*Δ to various stressors were identical for the parental strain and the mutant strain. They were respectively 0.062% for SDS, 50mM for DTT, >250 µg/mL for Cyclosporine >25 µg/mL for Caspofungin, >500 µg/mL for Nikkomycin, >25U for Zymolyase. The *kre6*Δ mutant was however more sensitive to the general cell wall inhibitors Congo Red and Calcofluor White than the parental strain (MICs for Congo Red: 50 µg/mL for *kre6*Δ and 500 µg/mL for the parental strain; MICs for Calcofluor White: 0.5 mg/mL for *kre6*Δ and 2.5 mg/mL for the parental strain (**Fig. 4**). These two compounds are sensitivity markers of all *kre6*Δ mutants

**Fig 4.**
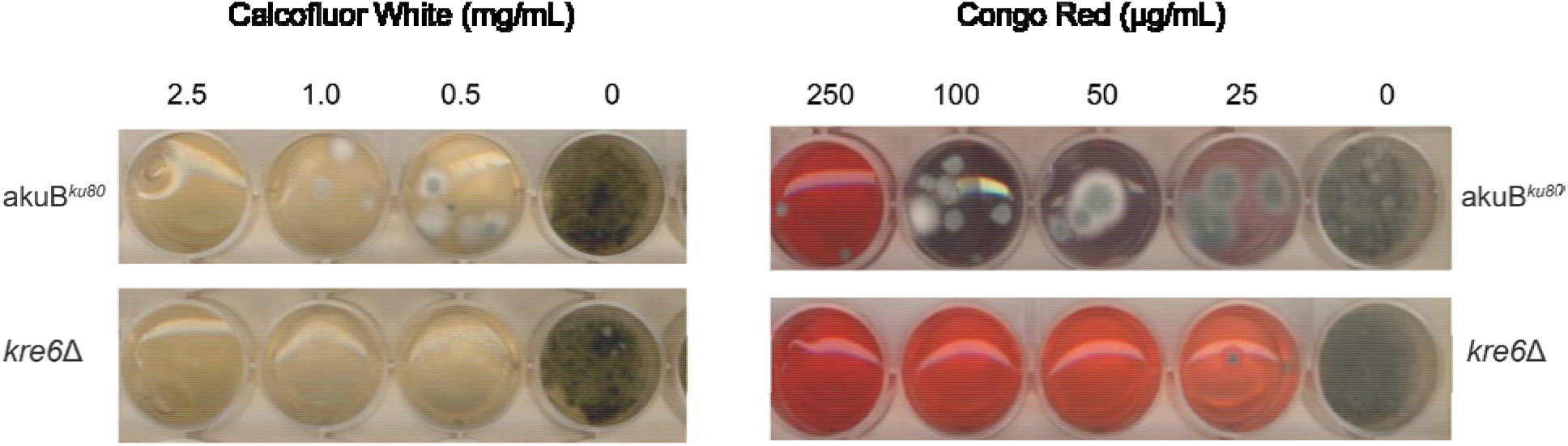
Sensitivity of *A. fumigatus* strains to Calcofluor White (left panel) and Congo Red (right panel). For each inhibitor, both the *akuB*^*ku80*^ (top) and *kre6*Δ (bottom) strains are tested.

## DISCUSSION

The structure and biological function of the β-1,6-glucan glucans are different in yeasts and mold. Solid-state NMR spectra revealed the presence of β-1,6-glucan signals. The sheared DP J-INADEQUATE spectra, which identify mobile components, demonstrated that β-1,6-glucans are the predominant component in the mobile phase of *C. albicans* (**Fig. 2**). In contrast, the detection of β-1,6-glucan signals at carbon positions C3, C4, and C6 in the spectra of *A. fumigatus* conidia indicates their presence in the rigid fraction of the conidia cell wall. In yeast, the β-1,6-glucans present in the mobile fraction is composed of 140-350 units (2). Recent structural data have shown that in *C. albicans* like in *S. cerevisiae*, β-1,6-glucans are composed of a main linear chain of β-1,6-glucosyl residues with side chains of β-1,3-glucosyl or β-1,3-laminaribiosyl residues (2). Even though the length and branching of the β-1,6-glucan in *A. fumigatus* is unknown, its affiliation to the rigid fraction in *A. fumigatus* would suggest it is formed as a long polymer.

The current study also confirmed that the synthesis of β-1,6-glucans is controlled by *KRE* genes in all fungi. The importance of the different *KRE* genes varies depending on the fungal species analyzed (21). Even though the exact enzymatic function of Kre6 proteins in *A. fumigatus* has not been characterized, it is clear that Kre6 proteins is responsible for the synthesis of this carbohydrate polymer in the cell wall of the conidia in *A. fumigatus*. However, the morphological role of this protein differs between different species. The *kre6*Δ mutants in *C. albicans, S. cerevisiae* and *C. neoformans* show similar phenotypes: partial defect in cell separation, hypersensitivity to Calcofluor white and Congo Red with a decreased amount of cell wall β-1,6- glucan in yeast (1, 4, 22). The phenotype of the *kre6*Δ mutant in *A. fumigatus* is very similar to the parental strain. The increased sensitivity of the *kre6*Δ mutant to well-known cell wall inhibitors such as Calcofluor White and Congo Red is the only similar phenotype seen in all *kre6*Δ mutants (**Fig. 4**). It suggests a role of β-1,6-glucan in the permeability to certain cell wall drugs. It has been indeed shown that the effect of Congo Red in *A. fumigatus* is dependent on the permeability of *A. fumigatus* to this drug (23). It is also known that *kre6*Δ mutants are not more sensitive to antifungal cell wall drugs such as caspofungin in yeasts (24). This is also confirmed for this Δ*kre6* mutant of *A. fumigatus*.

Strong signals for β-1,6-glucan are only detected in the conidia of *A. fumigatus akuB*^*ku80*^ cell wall while they are missing in *kre6*Δ mutants (**Fig. 3C**) and in the mycelium of the parental *akuB*^*ku80*^ (**Fig. 1A**). This is another example of the significant differences in the composition of the cell wall of the conidium and the mycelium of a filamentous fungus. In *A. fumigatus*, the outer layer of the conidium contains specifically melanin and rodlets. Moreover, a new specific constitutive polysaccharide was recently identified in the conidia (25). It is a specific mannan with unique linkages and with side chains containing galactopyranose, glucose, and N-acetyl glucosamine residues. The analysis of mutants lacking one to three major components of the outer layer of the conidial cell wall showed that the absence of one component of the outer layer of the conidial cell wall such as α◻1,3-glucan, melanin, and rodlet proteins led to unexpected phenotypes without any logical additivity. Unexpected compensatory cell wall surface modifications were indeed observed, such as the synthesis of the galactosaminogalactan, the increase in chitin and glycoprotein concentration or changes in permeability.

The increase in concentration of β-1,3/1,4-glucan in the *kre6*Δ mutant (**Fig. 3B, C**) indicated some compensatory reactions to the lack of β-1,6-glucan without any significant morphological changes. This result is not surprising since the deletion of the β-1,3/1,4-glucan synthase does not any impact on the *A. fumigatus* growth (26). In contrast, the main scaffold composed of the majority of the α-1,3-glucan, β-1,3-glucan and chitin are not affected by the loss of β-1,6-glucan. The putative role of β-1,6-glucan in *A. fumigatus* could be to indirectly stabilize the network of these major polysaccharides even though it is in very low concentration. This result also suggested that the loss of rigid molecules in low concentrations such as β-1,6-glucan glucans or galactomannan (20) does not lead to major changes in the conidia cell wall which exhibited similar presence of the major structural polysaccharides which are chitin, α-1,3-glucan and β-1,3- glucan.

## MATERIALS AND METHODS

### Preparation of fungal materials and labeling of *A. fumigatus* for solid-state NMR

The fungal strain *A. fumigatus akuB*^*ku80*^ was used (27). The strains *A. fumigatus akuB*^*ku80*^ and *kre6*Δ were cultured on agar plate containing 20g/L ^13^C-glucose (Catalog # CLM-1396-PK, Cambridge Isotope Laboratories) and sodium nitrate salt solution (NLM-712-PK, Cambridge Isotope Laboratories) as the sole carbon and nitrogen sources, respectively, supplemented with trace elements (**Table S5**) incubated at 37ºC for 3 days. Conidia were collected from the plate using aqueous 0.5% Tween 20 solution washed with 2 times of deionized water, followed by a phosphate-buffered saline (PBS) to remove excess salt and glucose and centrifugated at 3000 rpm for 10 min. The intact conidia were packed in 3.2 mm rotor for solid state NMR analysis.

High-resolution solid-state NMR experiments were conducted on a Bruker Avance Neo 800 MHz spectrometer equipped with a 3.2 mm HCN triple-resonance MAS probe, located at the Max T. Rogers NMR Facility at Michigan State University. For ^13^C detection, magic-angle spinning (MAS) was performed at a frequency of 13.5 kHz, and all experiments were carried out at a regulated temperature of 293 K. The ^13^C chemical shifts were externally referenced by calibrating the methylene (CH◻) resonance of adamantane to 38.48 ppm, which served as the secondary reference standard for all spectra acquired from fungal samples. Radiofrequency (RF) pulse field strengths ranged from 71.4 to 83.3 kHz for ^1^H hard pulses, decoupling, and cross-polarization (CP) transfers, while ^13^C pulses were applied with RF field strengths 50 and 62.5 kHz.

### Identification of carbon resonances corresponding to carbohydrate moieties

To elucidate the molecular architecture of the fungal cell wall, a series of two-dimensional solid-state NMR experiments were conducted. The through-bond ^13^C-^13^C connectivity’s of dynamically mobile constituents were characterized using the refocused J-INADEQUATE experiments (28, 29), performed under a direct polarization (DP) scheme with a recycle delay of 2 s (**Fig. S6**). The J-evolution period comprised four delays of 2.3 ms each, optimized to enhance the sensitivity toward carbohydrate moieties. In contrast, the structurally rigid components were investigated using the CORD (COmbined R2_n_ ^v^-Driven) sequence (10, 11), which employed CP for initial magnetization transfer and a mixing time of 53 ms under 13.5 kHz MAS conditions (**Fig. S3**). The resulting two-dimensional spectra enabled detailed resonance assignments of carbon sites within the constituent polysaccharides. These assignments were cross validated using reference chemical shift values from the Complex Carbohydrate Magnetic Resonance Database, as well as recent literature reports (6, 30-32). A comprehensive list of assigned chemical shifts is provided in **Table S1**. The experimental parameters are documented in **Table S6**.

### Compositional analysis of cell wall carbohydrates

The molar composition of carbohydrates in each sample was assessed through quantitative analysis of peak volumes obtained from two-dimensional ^13^C-^13^C correlation spectrum. These spectra were acquired using both the 53 ms CP- CORD and direct polarization (DP) refocused J-INADEQUATE experiments, which selectively probe the rigid and mobile polysaccharide fractions, respectively. Peak volumes were extracted using the integration module of Bruker Topspin software (version 4.1.4). To minimize uncertainties associated with spectral overlap, only well-resolved and unambiguous signals were selected for quantitative analysis. In the CORD spectra, quantification was performed by averaging the volumes of clearly resolved cross-peaks attributed to each polysaccharide. For the INADEQUATE spectra, only well-defined scalar-coupled carbon pairs were included in the analysis. The specific ^13^C NMR resonances utilized for compositional estimation, along with their chemical shift assignments and integrated intensities, are detailed in **Tables S2** and **S3**. Relative molar abundances were determined by normalizing the integrated peak volumes to the number of contributing resonances for each carbohydrate species. Standard errors were calculated by dividing the standard deviation of integrated volumes by the number of cross-peaks analyzed. The overall standard error for each sample was derived from the square root of the sum of squared individual errors, following previously established methodologies (33, 34).

### Construction of the *kre6*Δ mutant

The deletion of *KRE6* in the *A. fumigatus akuB*^*ku80*^ strain (27) was performed as described earlier using hygromycin B as a resistant marker (35, 36). Integration of the drug resistance gene at the correct locus for *KRE6* was confirmed by Southern blot and PCR analysis (**Table S4** and **Figs. S1, 2**). cDNA *KRE6* was obtained by PCR using the set of primers 5’Kre6pREPter and 3’Kre6pREPter and *A. fumigatus* total cDNA as previously described (37).

### Growth, sporulation, germination, viability and morphology of the *kre6*Δ mutant

Mycelial growth was tested on different agar media (Malt2% (Malt 2%, Roswell Park Memorial Institute medium (RPMI), minimal medium (MM), and Glucose 3%-Yeast Extract 1% (YG) at 37 °C and 50 °C and the diameter of the colony was measured at different time points (24 h, 48 h, 72 h, and 96 h) after inoculation of 3 µL of a conidia solution at 5 × 10^6^ conidia/mL (c/mL). For the liquid condition: 10^6^ conidia were inoculated in 50 mL of liquid medium (3% glucose + 1% yeast extract) at 37 °C, 150 rpm. To calculate the dry weight of the fungus at each time point (24 h, 48 h, and 72 h), the mycelium was recovered on a filter (Macherey Nagel^®^ (Hœrdt, France): 0.55 mm), washed with water, dried at 80 °C for 24 h, and weighed.

The conidiation rates were estimated following the inoculation of 10^5^ conidia onto three tubes containing 2% Malt agar (10 mL/tube). After one week of growth at 37 °C, conidia were recovered with 5 mL of an aqueous 0.05% Tween 20 solution, and their number was counted with an hemacytometer.

Conidia were collected from agar slants/plates after seven days of growth at room temperature using 0.05% Tween 80 solutions. Conidial germination was followed microscopically every 30min at 37 °C. The sensitivity to Congo Red and Calcofluor-White (CR; C6277 and CFW; F3543, Sigma, St Louis, MO, USA was estimated on minimal agar medium containing different concentration of drugs (CR: 0 to 300 µg/mL and CFW; 0 to 200 µg/mL). A total of 3 µL of a suspension of conidia at was inoculated on plates and incubated at 37 and growth was measured at different times.

## ACKNOWLEDGMENTS

The solid-state NMR analysis was supported by the National Institutes of Health (NIH) grant R01AI173270 to T.W.

