## Supplementary file for "Kre6-Dependent β-1,6-glucan Biosynthesis Only Occurs in the Conidium of *Aspergillus fumigatus*"

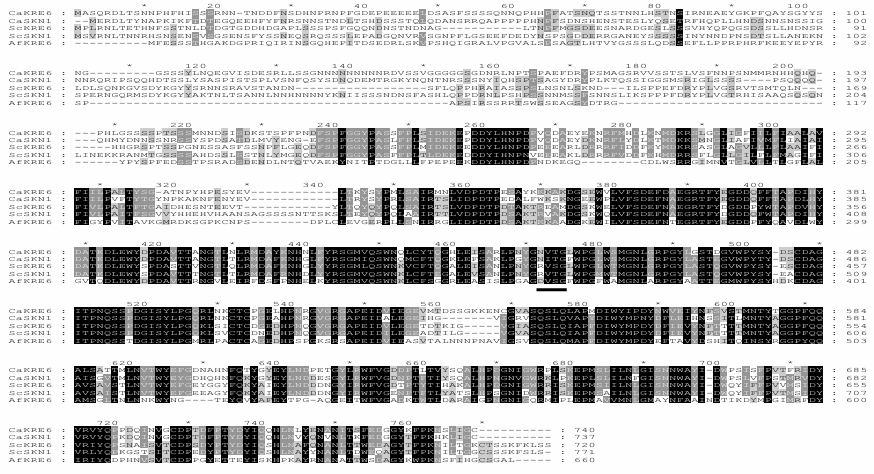

**Fig S1.** A multiple sequence Crustal X alignment of the *KRE6* and *SKNI* homologs in *C. albicans, S. cerevisiae* and *A. fumigatus.* Conserved amino -acids residues are in black. Similar amino acids residue is grey. Putative UDP glucose binding site is underlined.

**B**

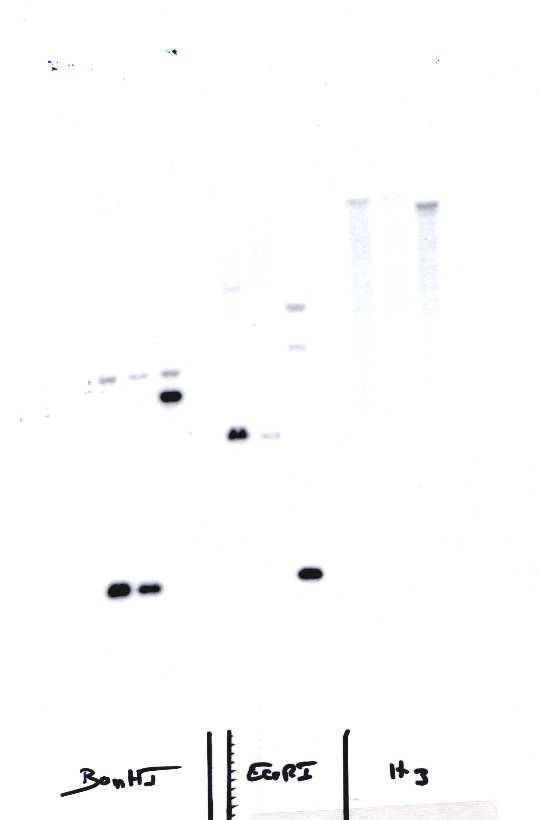

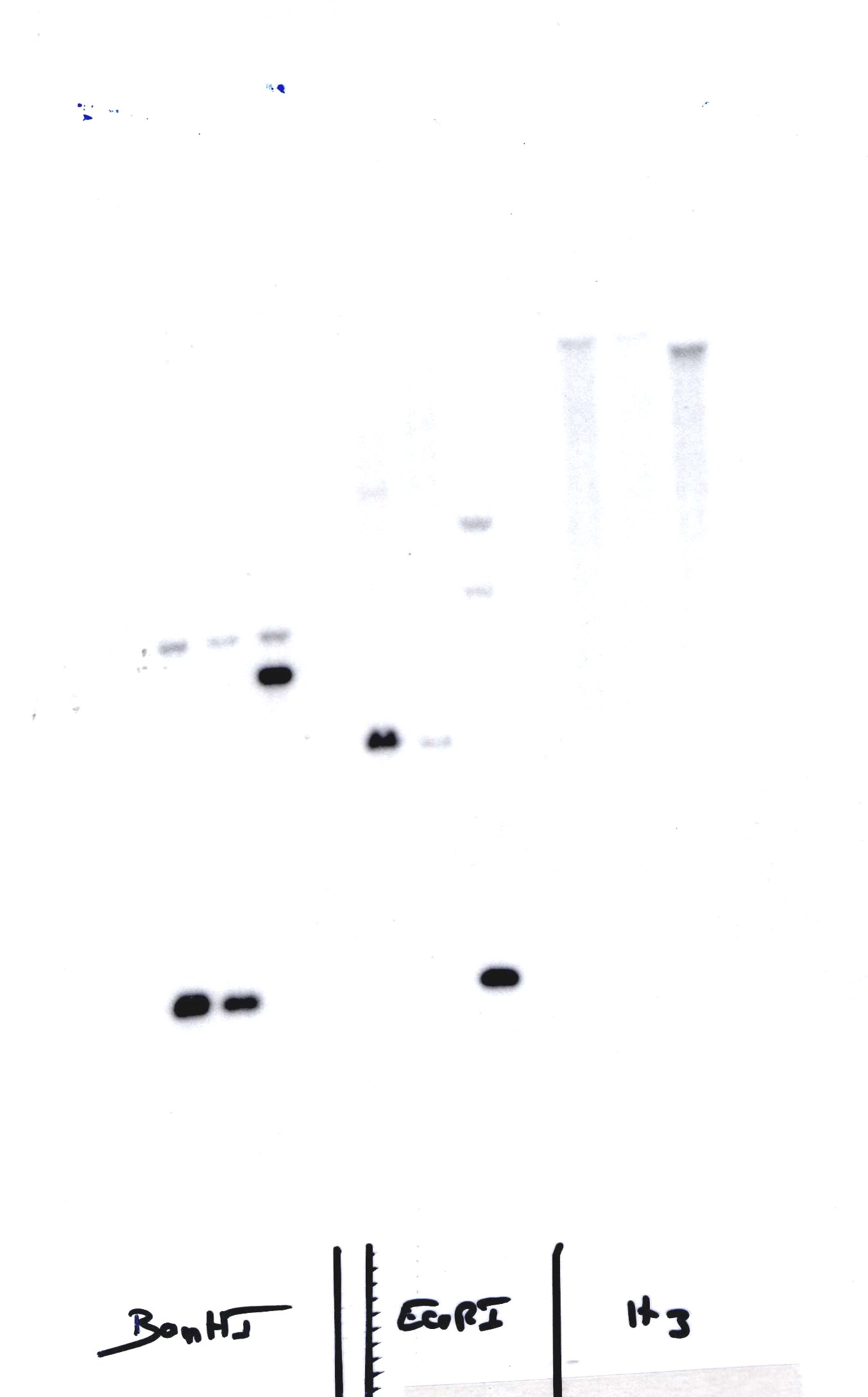

BamHI

EcoRI

1 2 3

1 2 3

2,3 Kb

1,3 Kb

4,0 Kb

3,0 Kb

1,5 Kb

**HindIII P2**

**pUC19-Δkre6-*HPH***

BamHI

BamHI

pUC 19

**BamHI P1**

StuI

XbaI

*KRE6*

*HPH*

*HPH*

pUC 19

*akuB^ku80^*

AfΔkre6

1 kb

BamHI

EcoRI

StuI

Xba I

HindIII

StuI

Xba I

HindIII

EcoRI

EcoRI

BamHI

probe

EcoRI

BamHI

**Fig. S2.** Characterization of *kre6*Δ mutant. (a) Schematic representation showing the predicted disruption events resulting from homologous integration of *KRE6* deletion construct into *A. fumigatus* genome. (b) Southern blot analysis of two *kre6*Δ mutants (lanes1-2) compared to the strain *akuB^ku80^* (lane 3). Genomic DNA was digested with BamHI or EcoRI and hybridised with labeled probe in (A).

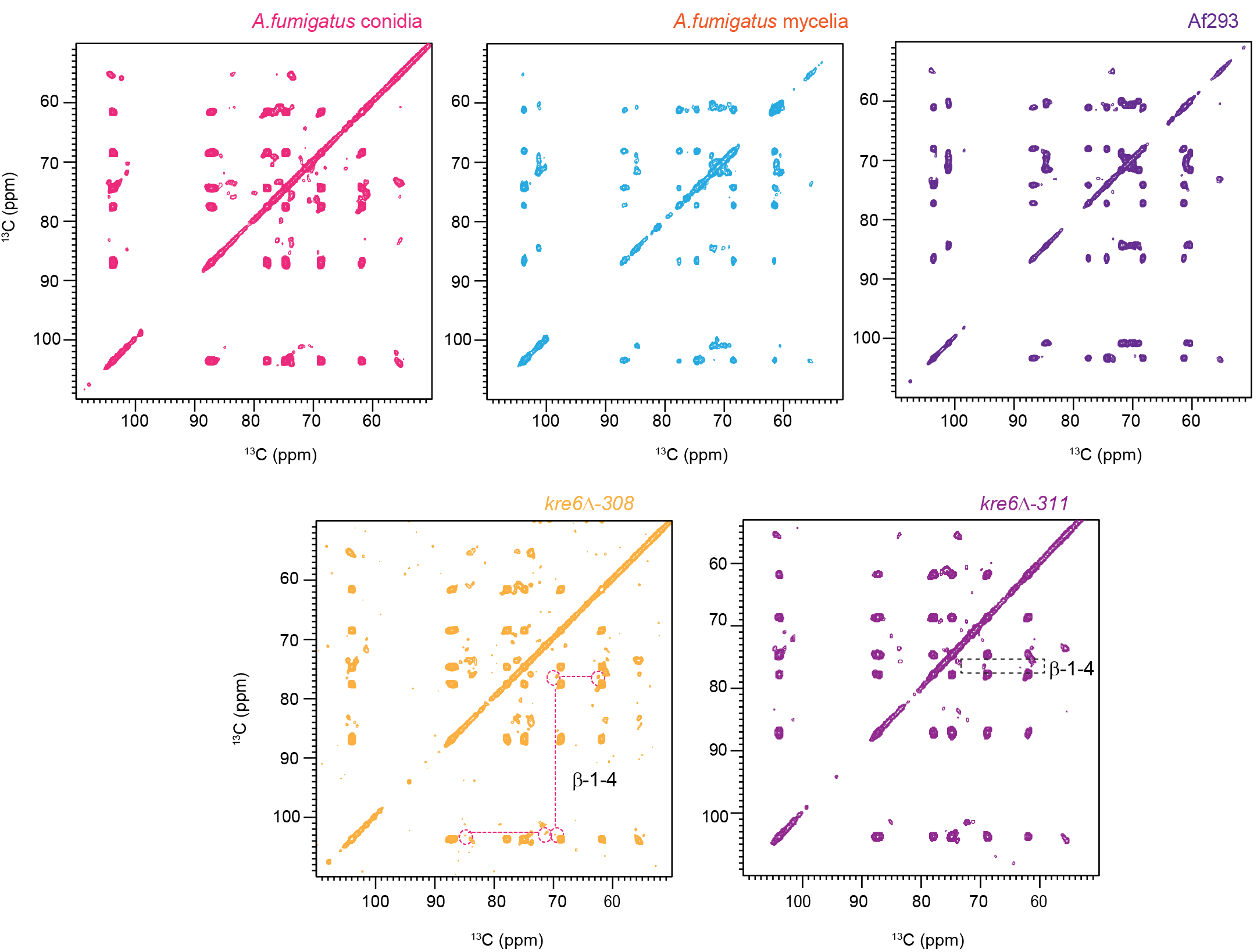

**Fig. S3.** Changes in rigid carbohydrates in *A. fumigatus* conidial cell wall *akuB^ku80^* (pink), mycelia(blue), Af293 *kre6*Δ mutants (*kre6*Δ-308 yellow) and (*kre6*Δ-311 magenta). The 2D ^13^C-^13^C CORD spectra of *kre6*Δ mutants (*kre6*Δ-308 yellow) and (*kre6*Δ-311 magenta) showing the new peaks for β-1,4-Glc unit in β-1,3/1,4-glucan. All spectra were recorded at 800MHz with 13.5 MAS.

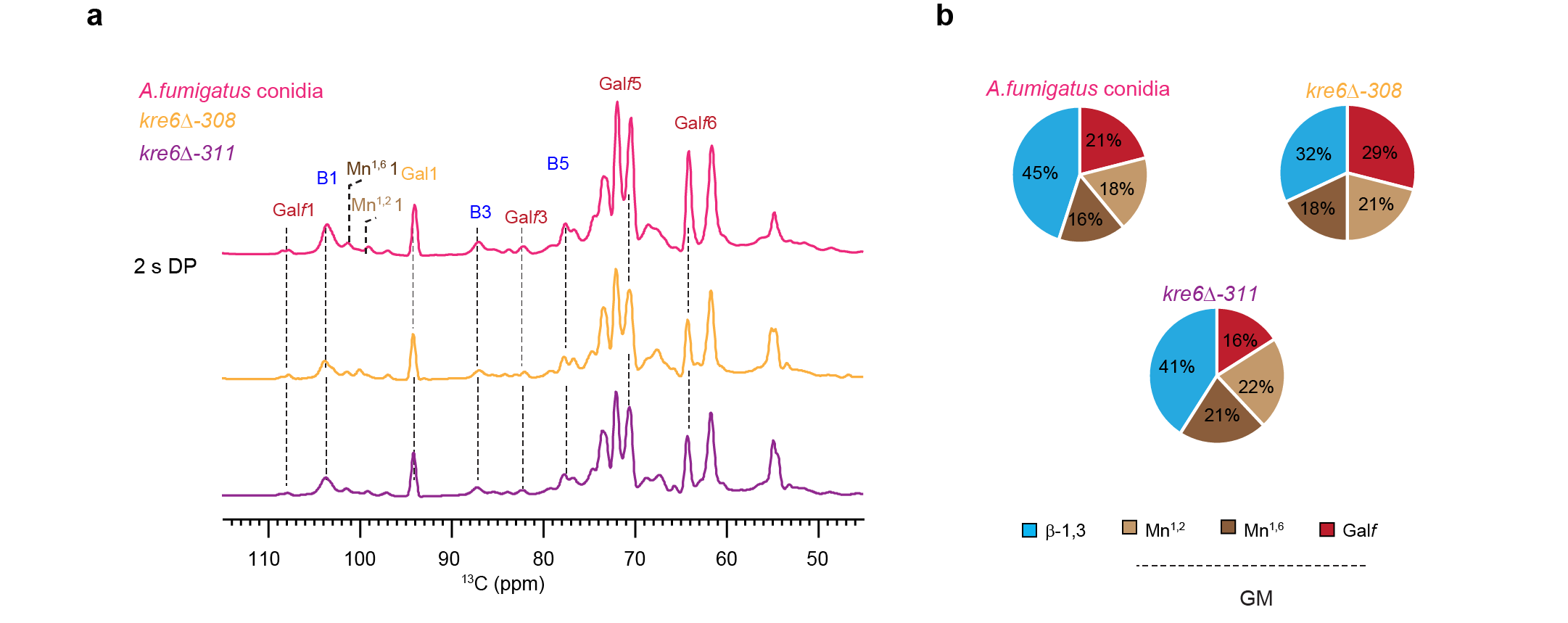

**Fig. S4. Mobile carbohydrate components in *A. fumigatus conidial* cell wall. (a)**  DP spectra with short recycle delay of 2 s for selection of mobile components has been shown for *A. fumigatus akuB^ku80^* (pink), *kre6*Δ mutants (*kre6*Δ-308, yellow) and (*kre6*Δ-311,magenta). **(b)** Molar composition of the mobile polysaccharides in *A. fumigatus* conidial cell wall, determined by peak volumes of ^13^C DP refocused *J*-INADEQUATE spectra.

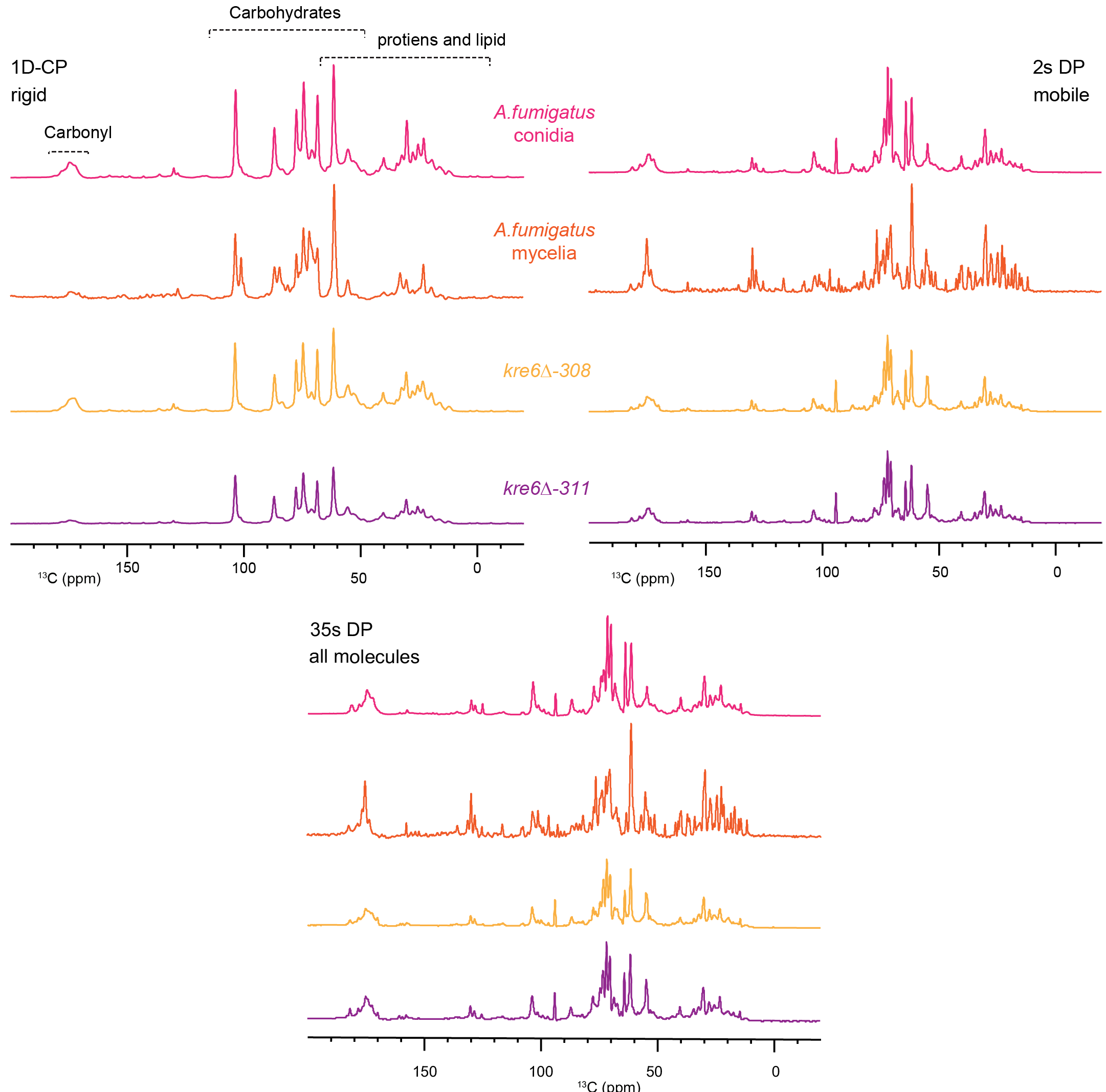
 **Fig. S5.** Proteins and lipids in *Aspergillus fumigatus* dorC are predominantly localized within the mobile phase. An series of 1D ^13^C spectra, each optimized to detects different components with distinct dynamics are compared for *A. fumigatus* conidia,mycelia and *kre6*Δ mutants. These spectra includes the 1D ^13^C CP spectra (rigid molecules), 1D ^13^C DP spectra measured with short recycle delays of 2s (preferencial detection of mobile molecules) along with long recyle delays of 35s (quantitaive detection). All the spectra were measured on 800MHz NMR under 13.5kHz MAS.

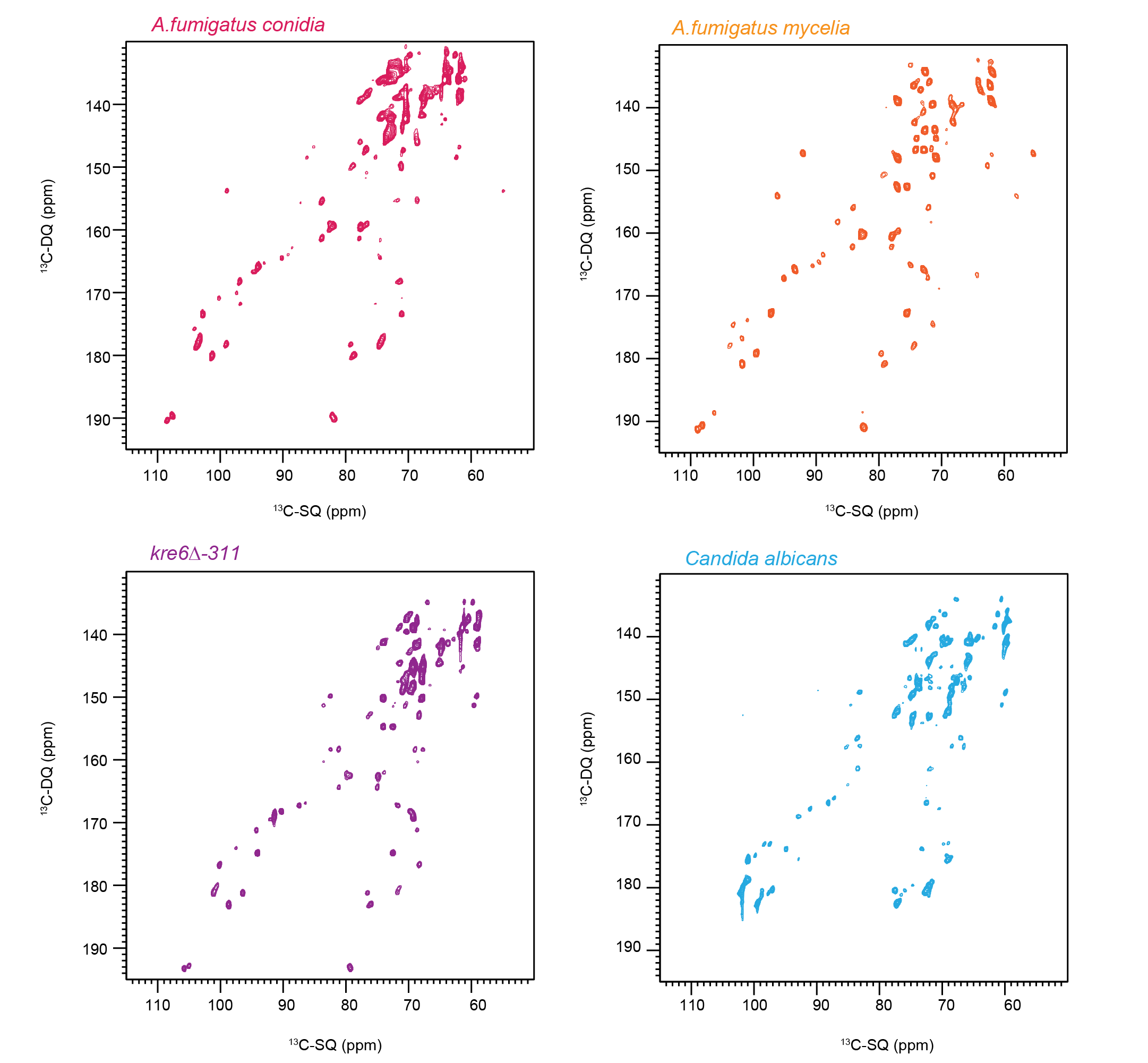

**Fig. S6** Changes in mobile carbohydrate region of *A. fumigatus* conidia(pink), mycelia(blue), *kre6*Δ mutants and *C. albicans.* 2D ^13^C DP *J*-INADEQUATE spectra of *A. fumigatus* (pink) showing carbohydrates region. 2D ^13^C DP *J*-INADEQUATE spectra of *C. albicans* showing the difference with *A. fumigatus* conidial cell wall.

**Table S1.**  ^13^C chemical shifts of *A*. *fumigatus* conidial cell wall. The referencing scale is TMS scale. Ambiguous sites are underlined: for Gl, the carbon numbering of C2, 3, 4 is uncertain.

| Carbohydrates | type | C1 | C2 | C3 | C4 | C5 | C6 | Reference |
| --- | --- | --- | --- | --- | --- | --- | --- | --- |
| Rigid molecules | | | | | | | | |
| β -1,3-glucan (B) |  | 103.6 | 74.4 | 86.8 | 68.4 | 77.6 | 61.4 | Shim *et al*. 2007^1^ |
| α-1,3-glucan (A) |  | 101.2 | 71.9 | 84.5 | 69.9 | 71.6 | 60.9 | Bhanja et al.2014^2^ |
| Chitin (Ch) |  | 103.9 | 55 | 73.2 | 83.3 | 75.5 | 61.5 | Fernando *et al.* 2021^3^ |
| Mannan |  | 102.3 | 71 | 63.9 | 67.1 |  | 61.9 |  |
| B1,6 glucan |  |  |  | 76.7 | 70.5 |  | 69.4 | Lowman *et al.* 2011 |
| Mobile molecules | | | | | | | | |
| β-1,3-glucan (B) |  |  |  |  |  |  |  | Shim *et al.* 2007  Fairweather *et al.* 2009  Saito *et al.* 1979 |
| β-1,5 galactofuranose (Gal*f*) | a | 108 | 81.8 | 77.5 | 83.5 | 71.7 | 63.5 | Chakraborty *et al.* 2021^4^ |
|  | b | 107.5 | 81.9 | 77.4 |  |  |  |  |
| α-1,6-Mannan (Mn^1,6^) |  | 102.7 | 70.8 | 73.8 | 67.7 | 70.8 | --- | Latge *et al.* 1994^5^  Chakraborty *et al.* 2021 |
| α-1,2-Mannan (Mn^1,2^) | a | 101.1 | 78.5 | 70.9 | 67.8 | 73.98 | 62.1 | Kuraoka *et al.* 2021^6^ |
|  | b | 98.9 | 79.0 | 71.19 | 67 | 73.4 | 61.98 |  |
| Galactose/Glucose or their derivatives  (Gl) | a | 90.04 | 74.0 | 74.5 | 70.27 | --- | 61.84 | Fontaine *et al.* 2011^7^ |
|  | b | 97.0 | 71.08 | 74.36 | 67.03 | --- | 61.82 | Archbald et al. 1981^8^ |
|  | c | 94.66 | 72.5 | 73.59 | 67.42 | --- | 61.9 | Archbald et al. 1981^8^  Fontaine *et al.* 2011^7^ |

**Table S2**. Molar composition of rigid polysaccharides in *A. fumigatus* dorC cell wall. The numbers were calculated using the integrals of well-resolved cross peaks of β-1,3 glucan and chitin in 2D ^13^C-^13^C CORD spectra. The results were already normalized by the number of scans.

| Strain | α-1,3-glucan | β-1,3-glucan | Chitin | β-1,6-glucan | β-1,6-glucan |
| --- | --- | --- | --- | --- | --- |
| KU80 | 3 ± 0.8% | 80 ± 12% | 9 ± 3% | 8 ± 2% | ND |
| *kre6*Δ-308 | 2 ± 0.9% | 83 ± 12% | 12 ± 3% | ND | 2 ± 0.2% |
| *kre6*Δ-311 | 4 ± 1% | 83 ± 11% | 10± 2% | ND | 2 ± 0.3% |

The area of the following well-resolved cross peaks 53 ms CORD spectra are used:

β-1,3: the average of C1-C2/3/4/5 and C3-C2/4/5/6.

β-1,6: the average of C3/5-C4 and C5-C6.

Chitin: the average of C1-2/4/5, C3-C2, C4-C2/3/5, and C5-C2.

Mannan: the average of C2-C5 and C2-C6.

**Table S3**. Molar composition of polysaccharides in *A. fumigatus* conidial cell wall. The numbers were calculated using the integrals of well-resolved cross peaks of β-1,3 glucan and chitin in ^13^C DP *J*-INADEQUATE spectra. The results were already normalized by the number of scans.

|  | β-1,3-glucan | Mn^1,2^ | Mn^1,6^ | Gal*f* |
| --- | --- | --- | --- | --- |
| KU80 | 32 ± 10% | 21 ± 4% | 18 ± 3% | 29 ± 8% |
| *kre6*Δ-308 | 45 ± 10% | 18 ± 12% | 16 ± 3% | 21 ± 8% |
| *kre6*Δ-311 | 41 ± 11% | 22 ± 6% | 21± 4% | 16 ± 4% |

The area of the following well-resolved cross peaks of 2D ^13^C-^13^C *J*- INADEQUATE spectra are used:

β-1,3 (a): the average of C1-C2, C2-C3 and C5-C6

Gal*f*: the average of C1-C2, C2-C3, C3-C4 and C4-C5

Mannan: the average of C1-C2, C2-C3 and C3-C4

**Table S4. Deletion of *KRE6* gene**

A 2kb *KRE6* fragment was obtained by PCR using *A. fumigatus* *akuB^ku80^* DNA as template and the set of primers BamH1-P1 and HindIII-P2 (Table1.). The fragment was cloned into pUC19 at BamH1 and HindIII restriction sites. The 0.7kb StuI-XbaI fragment of HPH resistance gene flanked by the GDP promoter and TRPC terminator, obtained behind from plasmid pAN7.1(Punt,Oliver, Dingemanse,Pouwels,van den Hondel,1987). Gene disruption at the correct locus was confirmed by Southern hybridization, after digestion of the genomic DNA by two different restriction enzymes BamHI and EcoRI and by PCR amplification using sets of primer Kre6dinewa and Kredinewb.

| BamHI P1 | CGCGGATCCGCTCCTCAACGCATGGCGC |
| --- | --- |
| HindIII P2 | CCCAAGCTTAACATCCGGTACGGATTCA |
| Kre6dinewa | GATGAATTCAATACTGAAGGTC |
| Kre6dinewb | GATTCTAGATTTGCCGGGACT |
| 5’Kre6pREPter | TCGCGGCCGCATGTTCGAAAGCTCCTCAACGCAT |
| 3’Kre6pREPter | GACTCGAGTAAAGCTCCTGAACACCCGTGTAT |

**Table S5**. Recipe of mineral-based solid medium. The pH is adjusted to 6.5 with H3PO4 or 0.25 M KOH. Each sample uses 100 mL of medium that contains 2g agar, 2 g of 13C- glucose and 5ml of sodium nitrate solution with 0.1ml of trace elements. The culture media and condition were adapted from a previously described protocol^9^

|  | Reagent | For 1L |
| --- | --- | --- |
| Trace elements | ZnSO_4_.7H_2_O | 22.0g |
|  | H_3_BO_3_ | 11.0g |
|  | MnCl_2_.4H_2_O | 5.0g |
|  | FeSO_4_.7H_2_O | 5.0g |
|  | CuSO_4_.5H_2_O | 1.6.0g |
|  | CoCl_2_.6H_2_O | 1.6.0g |
|  | (NH_4_)_6_Mo_7_O_24_.4H_2_O | 1.1.0g |
|  | EDTA | 50.0g |
| Nitrate salt solution | NaNO_3_ | 300.0g |
|  | KCl | 26.0g |
|  | MgSO_4_.7H_2_O | 24.0g |

**Table S6.** Solid-state NMR experiments and parameters. To be quantitative, direct pulse (DP) experiments with 35 s long recycling delay were used. cross polarization (CP), most rigid molecules. With DP and a shorter recycling delay of 2 seconds, suppress the rigid molecules from the spectra, and with Insensitive Nuclei Enhanced by Polarization Transfer (INEPT) the most mobile molecules were selected. For 2D 13C-13C correlation experiments allowed to resolve rigid intramolecular peaks. 2D DQ-SQ, DP J-INADEQAUTE and CP INADEQUATE spectra were used to detect through-bond correlations. The experimental parameters include the 1H Larmor frequency, total experiment time (t), recycle delay (d1), number of scans (NS), The number of points for the direct (td2) and indirect (td1) dimensions, the acquisition time of the direct dimension (aq2) and the evolution time of indirect dimension (aq1), spectral width (sw1 and sw2).

| Experiment | B_0_ (T) | t (h) | d1 (s | NS | td2 | td1 | aq2 (ms) | aq1 (ms) | sw2 (ppm) | sw1 (ppm) | ν_MAS(_kHz) |
| --- | --- | --- | --- | --- | --- | --- | --- | --- | --- | --- | --- |
| 1D ^13^C CP | 18.8 | 0.5 | 1.8 | 1024 | 3600 |  | 18 |  | 496.8 |  | 13.5 |
| 1D ^13^C DP | 18.8 | 0.5 | 2.0 | 256 | 3600 |  | 18 |  | 496.8 |  | 13.5 |
| 1D ^13^C DP | 18.8 | 2.5 | 35.0 | 256 | 3600 |  | 18 |  | 496.8 |  | 13.5 |
| 1D ^13^C refocused INEPT | 18.8 | 1.5 | 4.0 | 1024 | 3200 |  | 16 |  | 496.8 |  | 13.5 |
| 2D ^13^C-^13^C with CORD mixing | 18.8 | 11 | 2.0 | 32 | 2800 | 600 | 14 | 7.5 | 496.8 | 198.7 | 13.5 |
| 2D ^13^C-^13^C refocused DP J-INADEQUATE | 18.8 | 6 | 1.5 | 16 | 2600 | 1024 | 19 | 10 | 326.8 | 248.5 | 13.5 |
